# A SIMPLE pipeline for mapping point mutations

**DOI:** 10.1101/088591

**Authors:** Guy Wachsman, Jennifer L. Modliszewski, Manuel Valdes, Philip N. Benfey

## Abstract

A forward genetic screen is one of the best methods for revealing the regulatory functions of genes. In plants, this technique is highly efficient since it is relatively easy to grow and screen the phenotypes of hundreds or thousands of individuals. The cost-efficiency and ease of data production afforded by next-generation sequencing techniques have created new opportunities for mapping induced mutations. The principles of genetic mapping remain the same. However, the details have changed, which allows for rapid mapping of causal mutations. Current mapping tools are often not user-friendly and complicated or require extensive preparation steps. To simplify the process of mapping new mutations, we developed a pipeline that takes next generation sequencing fastq files as input, calls on several well-established and freely available genome-analysis tools, and outputs the most likely causal DNA change(s). The pipeline has been validated in Arabidopsis and can be readily applied to other species.

## Introduction

Identifying genetic mutations that underlie phenotypic changes is essential for understanding a wide variety of biological processes. A forward genetic screen is one of the most powerful tools for searching for such mutations. Spontaneous and induced mutations have been used to identify genes underlying aberrant phenotypes for over one hundred years (Morgan 1910). In the common case, a mutagen is used to generate a few thousand random mutations in the genome by physical (radiation; (Muller 1927)), chemical (EMS; (Lewis and Bacher 1968)) or biological (transposons; (McClintock 1950)) agents followed by a screen for the desired phenotype caused by one of the mutations. Once a plant with the desired phenotype is isolated, the researcher must identify the causal mutation. This is done by testing for an association between known genetic markers and the phenotype. Such genetic linkage (assessed by co-segregation) indicates that the causal mutation is located in the vicinity of the genetic marker. The introduction of next-generation sequencing (NGS) for mapping purposes has proven to be very promising, as it is possible to quickly identify a small number of potential causal SNPs. However, the currently available tools e.g., SNPtrack (Leshchiner et al. 2012), SHOREmap (Schneeberger et al. 2009) and NGM (Austin et al. 2011) are either inoperable (SNPtrack) or require coding knowledge. We have developed the SIMPLE tool (SImple Mapping PipeLinE), which operates on the input of the NGS fastq files generated from WT and mutant bulked DNA pools, and produces tables and a plot showing the most likely candidate genes and locations, respectively. The tool can be easily downloaded and executed with no prior bioinformatics knowledge and requires only a few simple preparatory steps to initiate. Once the program runs, the user accesses a table with the most likely candidate genes and a figure file that marks the location(s) of these candidates. Our pipeline has several advantages in comparison to other mapping tools. First, the entire process is user-friendly. It does not require any prior programming knowledge or NGS analysis skills. Second, it is “all-inclusive”; besides a few initial steps such as downloading the fastq files to the right folder and naming them, the user only needs to paste three lines into the terminal application to run the program. These steps are described in the README file. Third, the program can accept as input any paired-end or single-end fastq combination from the WT and mutant bulks. Fourth, this tool accepts, based on our experience, several types of segregating populations such as M2, M3, back-cross and map-cross. The project is currently hosted on GitHub and is available for download at https://github.com/wacguy/Simple.

## Results

### The concept behind the mapping tool

All methods that aim to map a causal DNA change implement a similar strategy. For bulk-segregant analysis, a segregating F2 population is divided into mutant phenotype and WT phenotype. DNA from each of the two bulked samples is sequenced by next-generation sequencing. For recessive mutations, the basic principle is to find the DNA changes that have only non-reference reads (i.e., locations that differ from the reference genome) in the mutant bulk and ~1:2 mutant to WT ratio of reads in the WT bulk (since the WT bulk is comprised of +/− and +/+ individuals in a 2:1 ratio). For dominant mutations, the concept is similar, but the non-segregating (with a reference linked locus) bulk would be the WT batch (+/+) whereas the mutant bulk is comprised of mixed genotypes of mutant individuals (+/− and −/−). Our pipeline is based on a short BASH script that calls BWA (Li and Durbin 2009), Samtools (Li et al. 2009), Picard (http://broadinstitute.github.io/picard), GATK (McKenna et al. 2010; DePristo et al. 2011; Van der Auwera et al. 2013), SnpEff (Cingolani et al. 2012) and R (R Core Team 2013) in order to generate two VCFs (variant call files) and a plot. The first VCF lists all candidate genes and the second one contains the entire SNP population found by the GATK HaplotypCaller tool. Generation of the candidate list is based on several criteria. First, and most importantly, the mutation has to segregate in the correct ratio. Additionally, we chose to focus on SNPs that are consistent with a specific set of criteria to identify the most likely causal mutation. However, these criteria can be readily changed by manipulating the simple.sh file. For example, we select only single nucleotide changes, since single nucleotide mutagens such as EMS are commonly used. We also select for mutations that have a significant effect on the protein rather than synonymous mutations or changes in intergenic regions. Figure 1a shows a flowchart of the pipeline concept.

**Figure 1.**
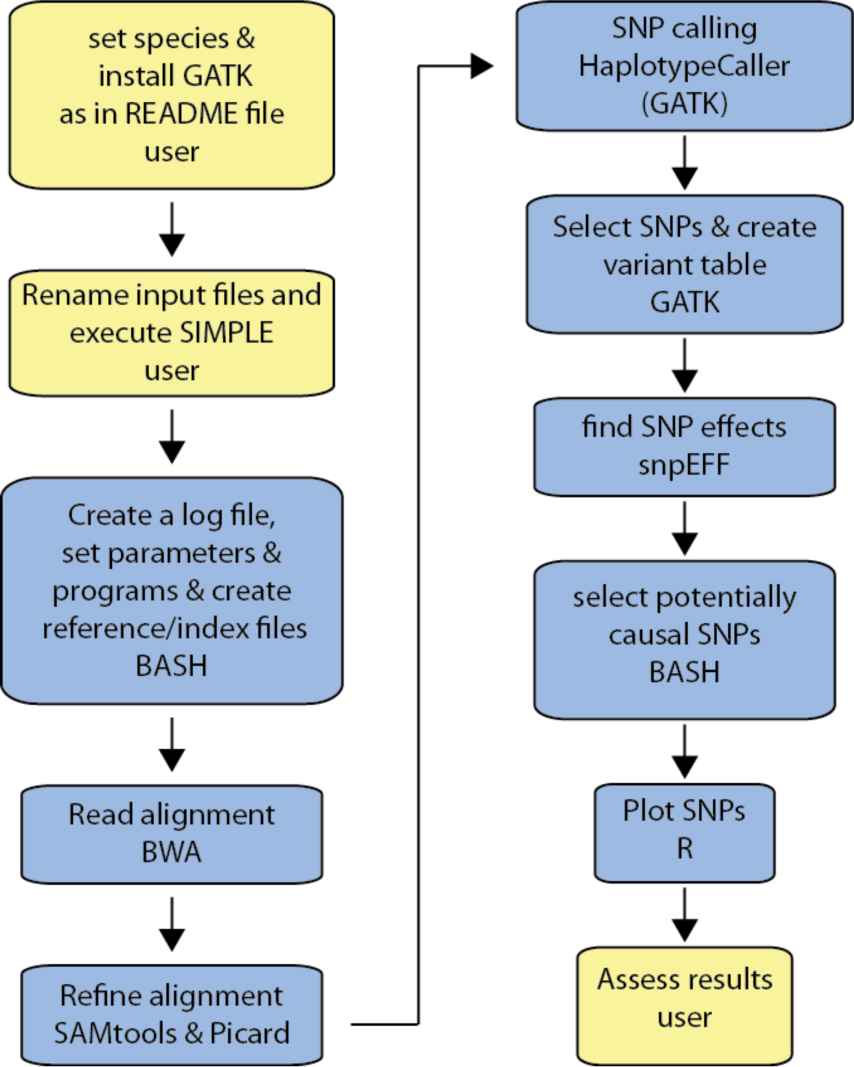
The SIMPLE pipeline workflow. User-required actions are in yellow and SIMPLE actions are in blue. The specific program that executes the action(s) of SIMPLE is indicated at the bottom of each blue box.

To test our pipeline, we sequenced ten populations that were generated in three independent EMS screens. We used four different population types including M2 (a segregating population generated by selfing a heterozygous plant from a mutagenized population), M3 (a segregating population generated by selfing a heterozygous M2 plant), back-cross (an F2 segregating population generated by crossing a mutant plant with the original parental line) and map-cross (an F2 segregating population generated by crossing a mutant plant with another accession, e.g., *L. erecta*). Some of the mutants were sequenced more than once or in successive generations, if they had more than one mapping population or if initial mapping failed (see Table 1 for details).

**Table 1.**
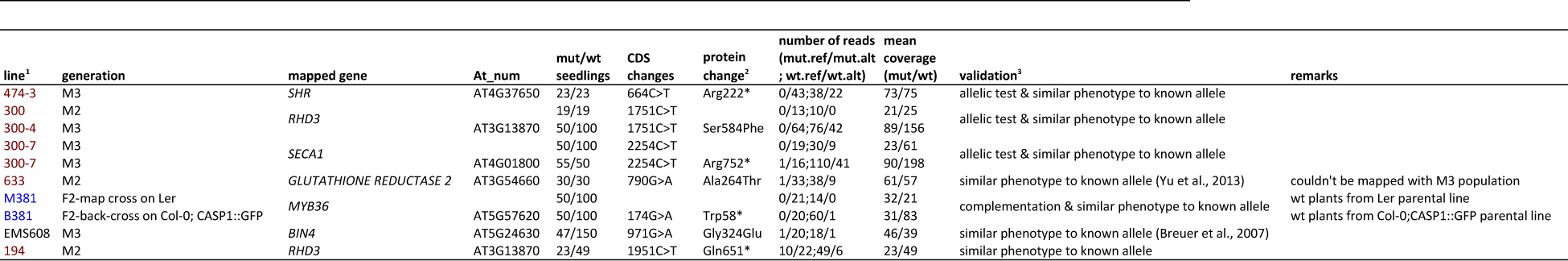
Mapping populations that were used to test the SIMPLE pipeline. Column 1 denotes the line number. Line 300-4 was generated by selfing a heterozygous plant from line 300. ^1^Each color represent a different EMS mutagenizing experiment; ^2^asterisk indicates a stop codon; ^3^see methods for more information.

The pipeline produces three files, cands4.txt, plot4.txt and Rplot.pdf (see details below) to help identify the causal mutation. In most cases, the candidate list output file (cands4.txt files) together with the position to SNP-ratio plot were sufficient to identify the causal mutation (Fig. 2a). In line 194 no genes were found in the candidate list, most likely due to an erroneous inclusion of WT seedlings in the mutant bulk. Nevertheless, the program provides additional information to help identify strong candidates. For example, the plot indicates a linked locus near the center of chromosome 3 (Fig. 2b). Browsing the SNP population in this region identifies a premature stop codon in *RHD3* (*AT3G13870*). Indeed, this line has a short root and wavy root hair phenotype similar to the mapped M2-300/M3-300-4 line and to *rhd3-1* (Wang et al. 1997; 2015).

**Figure 2.**
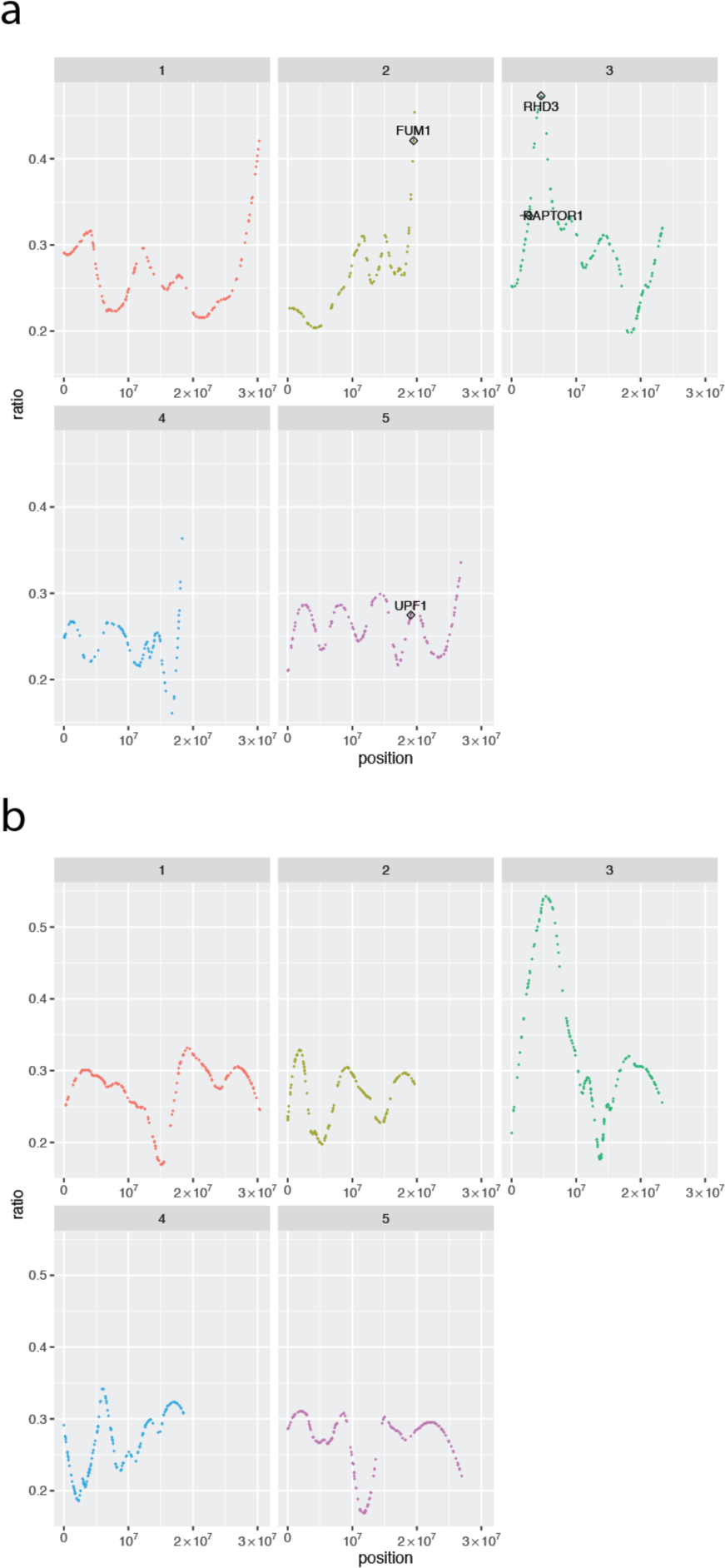
Example of the output plots. x-axis, chromosomal location; y-axis, ratio variable (see equation 1 for details). The number of each chromosome is labeled on top of each panel. a) and b) Line 300 and 194, respectively; see Table 1 and main text for details.

There are two parameters that appear to strongly influence the success of the pipeline in identifying the causal mutation or at least a short list of potential SNPs. First, incorrect inclusion of WT plants in the mutant bulk leads to reference reads in the mutant fastq file. As a result, the causal SNP and linked mutations are viewed as heterozygous. Therefore, it is essential that all individuals in the mutant bulk be correctly phenotyped as mutant. In cases where the phenotype is difficult to recognize, for example, as often happens with QTLs, it is recommended to work with smaller but high confidence populations rather than include potentially wild-type individuals in the mutant bulk. Alternatively, mutant plants can be tested based on the segregation of their offspring. A second crucial parameter is sequencing depth. This is important because different genomic regions have variable coverage depth and even when mean coverage appears to be sufficient, some regions may still have very few reads, which renders them almost impossible to genotype. We recommend a minimum of 30x coverage.

We recommend working with an F2/M2 generation rather than an F3/M3 generation for two reasons. First, a segregating F3/M3 population generated from a single F2/M2 heterozygous plant is homozygous for one quarter of the loci as well as being homozygous for the causal SNP whereas an F2/M2 population is homozygous only for the causal mutation and the genetically linked region. Therefore, there is substantial information loss in the F3/M3 generation and mapping is heavily dependent on the segregation of the WT bulk. This bulk is less informative because its 2:1 WT to mutant ratio in the binomial read-distribution has a higher potential to vary from the expected values. In contrast, the mutant bulk is expected to have strictly no reference reads and a few dozen alternate reads. Another reason to prefer the F2/M2 generation is that the following generation goes through a second round of recombination that generates chromosomes and chromosomal regions with more complex haplotypes that are more difficult to interpret.

#### Input files

The user provides the fastq files that are generated by a next-generation sequencing platform such as Illumina 2000 and places them in the fastq folder. Each bulk (mutant or WT) can have either a single file in the case of single-end sequencing or two files for paired-end. All other dependencies such as reference files and programs are either present or downloaded as part of the pipeline, with two exceptions. The user has to provide the GATK executable due to licensing issues; Java and R should also be pre-installed. A short description of how to run the pipeline is provided in the short README file (Supplementary file 3).

#### Output files

The program generates more than 30 files, although most of them are not necessary for non-programmers. There are three files that can help identify the causal mutation. The file Rplot.pdf (looks similar to Figure 2a or 2b) shows the chromosomal location of each SNP, plotted against a LOESS-fitted ratio variable. This variable was generated using this equation:

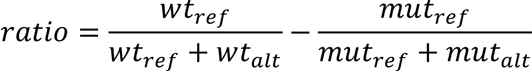

where,

*wt*_*ref*_ is the number of reads in the WT bulk that are called with the reference genome nucleotide,

*wt*_*alt*_ is the number of reads in the WT bulk that are called with a non-reference genome nucleotide,

*mut*_*ref*_ is the number of reads in the mutant bulk that are called with the reference genome nucleotide

and,

*mut*_*alt*_ is the number of reads in the mutant bulk that are called with a non-reference genome nucleotide.

The ratio should be around zero for unlinked SNPs and approximately 0.66 for the causal mutation and genetically linked SNPs (Figure 2). We removed all SNPs with a ratio lower than 0.1 and applied LOESS smoothing with degree=2 and span=0.3.

The second important file, cands4.txt lists the strongest candidate genes. (Supplementary file 1 as an example). These candidates are selected based on the following criteria. 1) The SNP has to be homozygous-alternate (namely, only reads that are different from the reference genome) in the mutant bulk and with approximately 2:1 reference to alternate read ration in the WT bulk. Second, the SNP must be a single nucleotide change rather than an indel. Third, the mutation should have a significant effect on the protein, i.e., splice acceptor, splice donor, start lost, stop gained or missense variant.

The third important file is plot4.txt (Supplementary file 2 as an example). This file lists all the SNPs with their locations, the change in the coding sequence, the effect on the protein and the number of reads for each allele in each bulk. This file is important in case the cands4.txt did not yield any candidate genes. For example, in some cases, the phenotype of the sampled mutants is difficult to distinguish from WT, such as the case when mapping QTLs or mutants with subtle phenotypes. In these scenarios, WT plants might be included in the mutant bulk and the causal SNP is interpreted as heterozygous for both bulks. In such cases, the user can manually browse the plot4.txt file to identify: 1) SNPs that are nearly homozygous-alternate in the mutant bulk and approximately 2:1 reference to alternate allelic ratio in the WT bulk and 2) SNPs with a significant effect on the coding region.

## Discussion

Approximately half of the Arabidopsis genes have unknown molecular functions (http://www.arabidopsis.org/portals/genAnnotation/genome_snapshot.jsp) which creates a large opportunity for discovering novel gene functionalities through relatively simple forward-genetic screens. The introduction of next-generation sequencing technologies offers new and rapid opportunities for identification of the genes affected by such screens. The main advantage of genome sequencing over traditional map-based cloning methods that use molecular markers such as RFLP and AFLP is that NGS provides single-nucleotide resolution. While pre-NGS methods could identify a genomic region, this had to be further mined for potential mutation(s) in candidate genes. By contrast, sequencing a mutagenized genome reveals the entire population of genetic changes. The causal mutation is then precisely identified using bioinformatics/computational tools. Another important benefit of using NGS for mapping is that a small population (a few dozen individuals) is usually sufficient for mapping. Since the mapping resolution is as high as a single nucleotide, the importance of each and every recombinant for reducing the region of the causal mutation (“chromosome walking”) is less important. In other words, the size of the region bounded by recombination events in which the causal mutation lies is not important as the researcher is able to visualize each and every SNP, evaluate it, and choose the one that has the highest likelihood of being causal. Mapping by NGS is especially fruitful when a large mapping population can be easily generated and screened which is the case for many plant species, nematodes and yeast. We have developed an easy to use tool that allows mapping of single-nucleotide induced mutations, even by researchers that have very little experience with bioinformatics tools. Our pipeline will take the fasq files as input and will identify causal SNPs with no pre-processing steps required. The output tables and plots can be readily used to identify the most likely mutation. Even in the case where no candidate gene is present, the list of SNPs with their read calls number in the mutant and WT bulks and their effect, will most likely point towards the correct gene. In theory, the SIMPLE pipeline can be used with any diploid species that has bulked mutant and WT mapping populations. Although not many species are currently covered, it is easy to add any species of interest to the pipeline (see README file). The program runs on Mac OSX version 10.11.6 and Linux release 6.7 (GNOME 2.28.2) with Java 1.7 installed (see README file for specification) which is a commonly used platform in many labs. The SIMPLE project is hosted on GitHub (https://github.com/wacguy/Simple) and includes a quick-start README file.

## Methods

### Growth condition and screening

Arabidopsis seedlings were grown on MS medium ((Murashige and Skoog 1962) containing 1% sucrose. EMS screen was performed according to (Weigel and Glazebrook 2006). DR5::Luc seedling were screened using the Lumazone system according to (Moreno-Risueno et al. 2010) with lines 474-3, 300, 300-4, 300-7, 633 and 194, with pCASP2::GFP expression for lines B381 and M381 (Liberman et al. 2015) and for the cortex marker (Brady et al. 2007) expression in the EMS608 line. The GFP screens for CASP2 and the cortex marker were performed using Leica fluorescence stereo dissecting scope.

### Sequencing

Paired or single-end libraries were prepared using NexteraXT or KAPA Hyper-prep Kit according to manufacturer instruction and sequenced on Illumina 2000/2500 instrument with the High or Rapid throughput mode at the Duke Center for Genomic and Computational biology.

### Mutant validation

Line 474-3 has a single ground tissue layer and a short root phenotype, similar to other *shr* alleles (Helariutta et al. 2000). The phenotype of F1 offspring in a cross between 474-3 as a male donor and *shr-2* (Helariutta et al. 2000) has the phenotype of the parental lines suggesting that line 474-3 is allelic to *SHR*.

Line 300 (and 300-4, a segregating line of 300) was crossed with *rhd3-1* (Wang et al. 1997; 2015) and the F1 progenies have short root and wavy phenotype similar to the parental lines suggesting that these lines are allelic. Line 300-7 was crossed with SALK line 063371 (Liu et al. 2010) and the F1 progenies phenocopied both parental lines. *myb36-1* (line 381 M and B) was previously described (Liberman et al. 2015). Line 194 has the same phenotype as line 300-4 and *rhd3-1* allele as described above.

All F1 crosses were sequenced for the male donor allele (see Table 2 for primers).

**Table 2.**
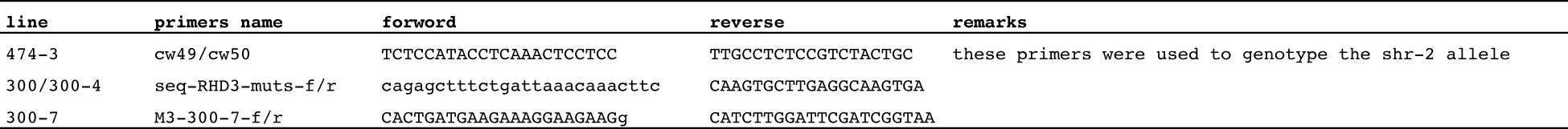
Primers that were used to amplify and sequence the male allele in F1 seedlings.

### Data access

Whole genome sequencing data have been deposited in the Sequence Read Archive (SRA) with the accession number PRJNA353239.

## Acknowledgments

We would like to thank people in the Benfey lab for comments on the manuscript and helpful experimental suggestions and to Carmen Wilson for technical assistant. We also thank the Duke Center for Genomic and Computational Biology for preparing and sequencing the DNA libraries. This research was funded by a grant to PNB from the NIH (R01-GM043778), and by the Howard Hughes Medical Institute and the Gordon and Betty Moore Foundation (GBMF3405).

## Supplementary file1

An example of cands4.txt file from line 633.

## Supplementary file2

An example of plot4.txt file from line 633.

## Supplementary file3

Instruction file for running the pipeline. This file is also available on the SIMPLE GitHub repository.

